# Multivariate comparison of taxonomic, chemical and technical data from 80 full-scale anaerobic digester-related systems

**DOI:** 10.1101/2023.09.08.556802

**Authors:** Pascal Otto, Roser Puchol-Royo, Asier Ortega-Legarreta, Kristie Tanner, Jeroen Tideman, Sjoerd-Jan de Vries, Javier Pascual, Manuel Porcar, Adriel Latorre-Perez, Christian Abendroth

**Affiliations:** Technische Universität Dresden, Institute of Waste Management and Circular Economy, Pirna, Germany; Darwin Bioprospecting Excellence, S.L. Parc Cientific Universitat de Valencia, Paterna, Valen-cia, Spain; Bioclear earth B.V., Groningen, the Netherlands; Institute for Integrative Systems Biology I2SysBio (University of Valencia - CSIC), Paterna, Spain; Brandenburgische Technische Universität Cottbus-Senftenberg, Chair of Circular Economy, Cottbus, Germany

**Keywords:** microbiome, 16S rRNA sequencing, anaerobic digestion, core microbiome, biogas, pa-rameter dependency

## Abstract

This study represents one of the most comprehensive characterisations of the anaerobic digestion (AD) microbiome with 80 samples from 45 different large-scale reactors in three countries. Technical, chemical and taxonomic data was thoroughly collected, analyzed and correlated to identify the main drivers of AD processes. Our results showed that MBA03, *Proteiniphilum*, a member of *Dethiobacteraceae*, and *Caldicoprobacter* were present in all the samples, while *Methanosarcina* was the most abundant and prevalent archaea. Two distinct bacterial clusters were identified by correlating microbial abundances. One was correlated with hydrogenotrophic and the other with acetoclastic methanogenesis. Organic acids, ammonia, nitrogen, COD and the trace elements Fe, Mo, and the macro nutrient P had the greatest impact on AD microbiomes. Temperature, reactor type and substrate also influenced the formation of a specialized microbial community. Overall, this work sheds light on the microbial key players involved in AD and evaluates how they are affected by technical and chemical parameters.

**Highlights:**

- Generation of a holistic dataset of chemical, taxonomic and technical parameters of 80 large-scale anaerobic digestion systems.
- Identification of a core microbiome comprising MBA03, *Proteiniphilum*, an uncultured organism from the *Dethiobacteraceae* family, and the *Caldicoprobacter* family.
- Identification of the main influencing parameters that determine the occurrence and non-occurrence of specific genera.
- Correlation of bacterial taxa with hydrogenotrophic and/or acetoclastic archaea.

## 1 Introduction

The production of biogas through anaerobic digestion (AD) can contribute to achieving key sustainability objectives such as the development of a circular economy, the transition to a bio-economy and the independent production of renewable electricity and gas (Golberg et al., 2016). Consequently, secondary raw materials such as organic waste, municipal sewage sludge, green waste, animal excrements, and agricultural residues can be converted into valuable resources such as methane and fertilizer. Compared to other forms of bioenergy production, the production of biogas through AD offers considerable advantages, as it closes nutrient cycles, it is the most energy-efficient process and it is the most environmentally friendly technology for the production of bioenergy from organic residues, linked to reduced emissions of climate-damaging gases such as methane and nitrogen oxides (Weiland, 2010). The conversion of these various organic materials into biogas is essentially carried out by complex microbial communities, that constitute the AD microbiome (Hassa et al., 2018; Q. Zhang et al., 2016).

AD is divided into four stages, each of which is carried out by different consortia of microorgan-isms with different environmental requirements (Weiland, 2010). In the first phase, hydrolysis, complex organic materials such as proteins, fats, and carbohydrates are hydrolyzed into the corresponding monomers and oligomers. These are then metabolized in the second phase, acidogenesis, to intermediates such as propionate, butyrate, other short-chain volatile fatty acids (VFAs), and alcohols. In the third phase, known as acetogenesis, acetic acid, CO_2_ and H_2_ are obtained by fermenting VFAs. Subsequently, these products are converted to CH_4_ and CO_2_ in a final phase, methanogenesis (Hassa et al., 2018; Weiland, 2010). The microbial communities of these four phases are generally categorized in fermentative bacteria, methanogenic archaea, and syntrophic bacteria. Fermentative communities carry out the first three phases, while methanogenic archaea use the intermediate products of the third phase to produce CH_4_ and CO_2_ in the fourth phase. Syntrophic communities are composed of microorganisms that work together in a mutualistic relationship (Leng et al., 2018). All those categories of microbial communities offer the opportunity to control and engineer the AD at different stages of the process.

The added value of investigating and understanding the microbiomes is that the efficiency, speed, robustness and adaptability of AD systems can be increased (Ni et al., 2020; Vrieze et al., 2016). However, microbiomes are very complex and depend on many factors such as pH, temperature, nutrients, etc., making it difficult to identify the microorganisms that are the main contributors to the efficiency of the process. Culture-independent methods such as 16S ribosomal RNA gene or metagenomics sequencing assist in the identification of crucial microorganisms and open new avenues for the complete characterization of the microbiome (McBee, 1954; Söhngen et al., 2014). Analysis of 16S rRNA gene amplicons has been used in several studies to comprehensively investigate microbial communities in AD systems (Perman et al., 2022; St-Pierre & Wright, 2014). Correlations between microbial profiles and various technical or physicochemical operating parameters such as substrates, organic loading rate (OLR), temperature, hydraulic retention time (HRT), etc. were established to assign changes in microbial community structure. In particular, studies on the core microbial communities are promising and help to identify important microbial taxa in running digesters. Their occurrence and abundance contribute to the understanding of the ecology of AD systems and can be linked to their function in the complete process as well as their relationship with relevant operating parameters (Kirkegaard et al., 2017; R. Xu et al., 2018).

Although multiple studies have already provided insight into the complexity of the underlying microbial communities, it is not yet clear how many microorganisms are involved in AD, what tasks each microorganism performs in the process, what this depends on, and whether there is a robust core microbiome. Puig-Castellví et al. found 1145 OTUs in a reactor that was co-digesting wastewater sludge (Puig-Castellví et al., 2020). Calusinska et al. observed 20 industrial biogas plants, and confirmed the hypothesis that different systems occur with different core microbiomes (Calusinska et al., 2018). Kirkegaard et al. compared nine full-scale digesters and looked for microbes present in all of them, which allowed the authors to narrow the core microbiome in these digesters down to 300 species. Moreover, these 300 OTUs explained 80% of all reads from all digesters (Kirkegaard et al., 2017). The presence of core microorganisms is strongly affected by the large number of influencing parameters. Researchers are increasingly using multivariate analyses to compare the large number of operational and physicochemical parameters with large DNA-based data sets. A particularly comprehensive study on this topic was recently conducted by Hassa et al. who compared 67 full-scale digesters from 49 agricultural plants (Hassa et al., 2021). In summary, Hassa et al. demonstrated the presence of microbial indicators for the specific process conditions analyzed (temperature, ammonia, substrate selection, etc.).

The present study contributes to understanding the microbial ecology of anaerobic digesters and their interaction with the operating conditions of the facilities, by providing one of the largest datasets to date on taxonomic profiles in AD systems with 45 different industrial digestion systems sampled in Germany, the Netherlands and Austria. It offers a complete and holistic dataset of microbial, chemical, and technical data from widely varying AD systems. Our findings expand the basic understanding of the AD microbiome, highlighting the microbial key players in the process and analyzing how different variables affect the underlying microbial communities.

## 2 Materials and Methods

### 2.1 Collection of the samples

A comprehensive and diverse set of 80 samples was collected from 45 full-size anaerobic digester plants. These reactors have been specially selected to represent a variety of different reactor systems, feedstock compositions and operational conditions such as temperature, HRT and OLR. Sampling took place from August to December 2021. A total of 80 distinct digester samples were collected in Germany, Austria and the Netherlands (Figure 1A). The Technical University of Dresden collected the samples in Germany and Austria, whereas the samples from the Netherlands were collected by Bioclear Earth B.V. In both cases, the sampling procedure was based on the same protocol. Samples for chemical analysis were collected in sterile 1 L sampling bottles. Samples for metagenomic DNA extraction and sequencing were collected in sterile 50-100 ml Falcon tubes. To preserve DNA, samples were mixed with pure ethanol directly upon collection. For this, a 1:1 ratio of sludge to ethanol was used. Samples for chemical analysis and for DNA sequencing were taken in triplicates. The collected samples were stored at -15 °C to avoid changes in the microbiome composition.

**Figure 1:**
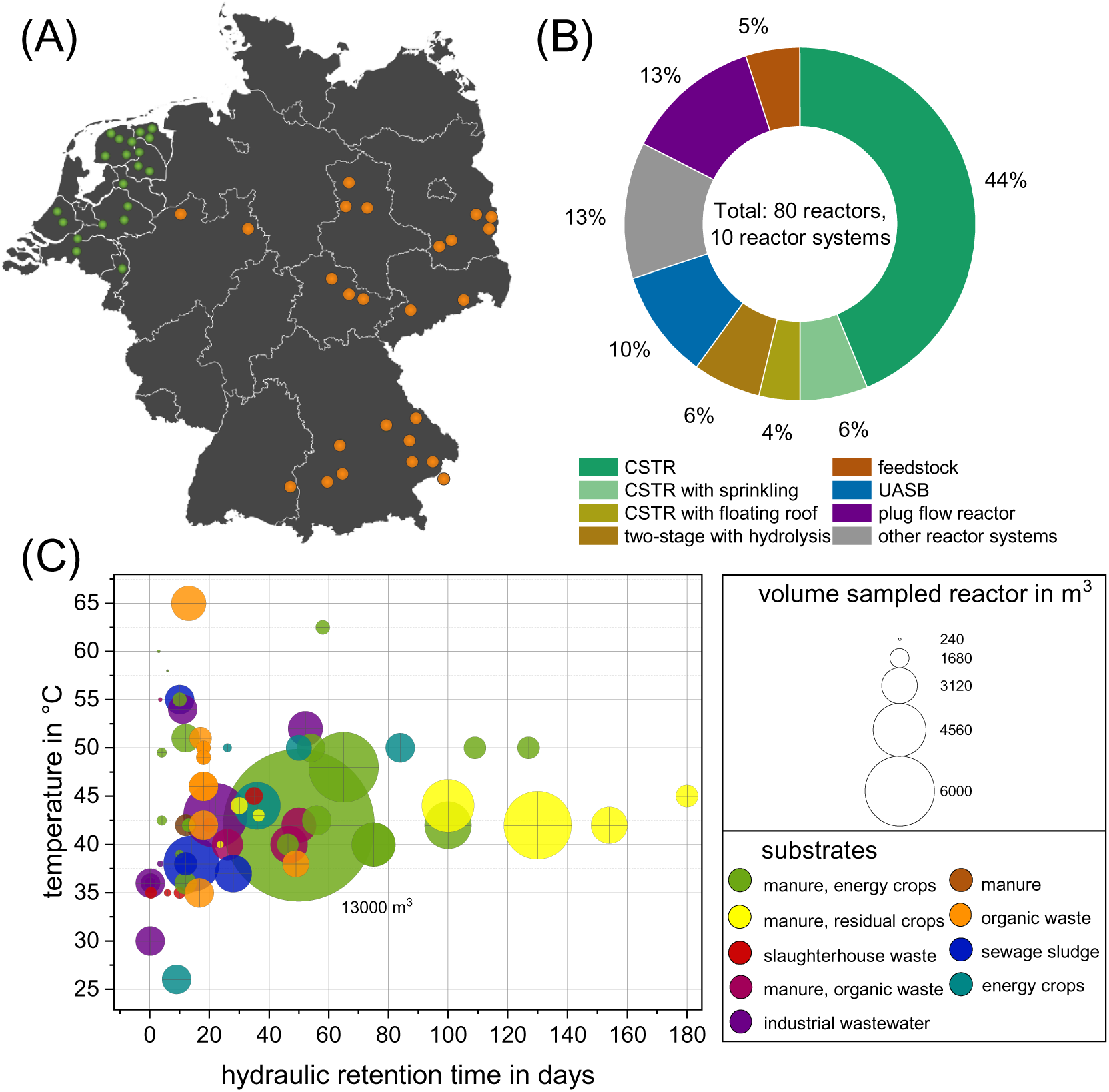
Overview of all sampled anaerobic digester systems: (A) Location of anaerobic digestion systems sampled in Germany (orange) and the Netherlands (green). (B) Types of anaerobic digester systems sampled. (C) Main operational data: temperature (y-axis) and the average retention time (x-axis) are shown. The bubble size indicates the reactor size and the colors indicate the main substrates in the respective feedstock.

### 2.2 Chemical and Technical data

Technical and operative data for each biogas plant were provided by the respective plant operator, including the operating temperature, reactor volume, hydraulic retention time (HRT), reactor type, substrate type, substrate quantity, years of operation, organic loading rate (OLR), additives, biogas yield, methane content, energy production and usage. An overview of the collected data is shown in Supplementary Table 1.

During the sampling phase, some parameters were directly measured on-site, for example, pH, conductivity, and the redox potential, which are sensitive to long storage times or regular freezing and thawing. The remaining chemical parameters were measured at the Technical University of Dresden as well as at Bioclear Earth B.V. These parameters include total solids (TS), organic total solids (oTS), chemical oxygen demand (COD), ammonium, total Kjeldahl nitrogen (TKN), FOS/TAC, the individual and total organic acids, heavy metals, and trace elements. All these parameters were determined using the appropriate norm (VDI-Gesellschaft Energie und Umwelt, 2016) and the raw data for the chemical parameters are shown in Supplementary Table 2. The concentration of the free ammonia (NH_3_) was calculated according to Hansen et al. including total nitrogen, pH and temperature (Emerson et al., 1975). The methods used to analyse the parameters and the corresponding instruments and units are shown in Supplementary Table 3.

### 2.3 DNA extraction and high throughput sequencing

An aliquot of 3 mL of each sample, conserved in ethanol, was centrifuged and washed with sterile Phosphate Buffered Saline (PBS) at least 3 times or until the supernatant was clear. DNA was extracted from the resulting pellets using the NucleoMag DNA Kit (Macherey-Nagel, Allentown, PA, USA) with the aid of the AutoPure96 robot for the purification step, following manufacturer’s instructions. DNA was quantified using the Qubit 1x dsDNA (Thermo Fisher Scientific, Waltham, Massachusetts, USA), and samples were sequenced by Novogene (Cambridge, UK). The extracted metagenomic DNA was used to amplify the hypervariable region V3-V4 of the 16S ribosomal RNA gene. The conserved regions V3 and V4 (470 bp) of the 16S rRNA gene were amplified using the following PCR cycle: initial denaturation at 95 °C for 3 min; 25 cycles of amplification (30 s at 95 °C, 30 s at 55 °C, 30 s at 72 °C); and 5 min of extension at 72 °C (Satari et al., 2020). The following primers were used: 341F (5’ CCTAYGGGRBGCASCAG 3’) and 806R (5’ GGACTAC-NNGGGTATCTAAT 3’). The amplification was carried out using the KAPA HiFi HotStart ReadyMix PCR kit (KK2602). The 16S rRNA amplicons were mixed with Illumina sequencing bar-coded adaptors (Nextera XT index kit v2, FC-131-2001), and libraries were normalized and merged. The pools with indexed amplicons were loaded onto the MiSeq reagent cartridge v3 (MS-102-3003) and spiked with 10 % PhiX control to improve the sequencing quality. Sequencing was conducted using paired-end 2X250pb or 2 × 300pb cycle runs on an Illumina MiSeq device.

### 2.4 Metagenomics and meta-data analysis

The raw Illumina sequences were loaded into Qiime2 (v. 2021.2.0) (Bolyen et al., 2019). The quality of the sequences was checked using the plugin Demux and the Qiime2-integrated DADA2 pipeline was used for trimming and joining the sequences, removing chimeras and detecting amplicon sequence variants (ASVs) (> 99.9% of similarity). The taxonomy of each sequence variant was determined via the classify-Sklearn module from the feature-classifier plugin, employing SILVA (v. 138) (Quast et al., 2013) as reference databases for taxonomic assignment. Microbiome data was analysed with the phyloseq package (v 3.16) in R (v 4.2.3) (McMurdie & Holmes, 2013). The abundance of each taxon was correlated to the abundance of the rest of microorganisms. Moreover, correlations between the microbial community and metadata were calculated. Due to the high number of taxa detected, the statistical tests were performed over the 250 most abundant microorganisms. Spearman’s rank correlation was used to study quantitative variables (i.e., content of nitrogen, concentration of acetic acid, etc.) and significant correlations were plotted into heatmaps using the Pheatmap package (v 1.0.12). Qualitative variables (i.e., mesophilic vs. thermophilic) were analyzed using the differential abundance test DESeq2 (v 3.16) (Love et al., 2014). Core microbiomes were calculated with the “coremicrobiome” function from the microbiome package (Lahti et al., 2017) in R (v 4.2.3).

## 3 Results and Discussion

### 3.1 Operational process parameters of the anaerobic digestion systems

80 full-scale anaerobic digesters and related systems were investigated based on chemical analyses and DNA sequencing as described in material and methods. Samples were collected at 45 sites in the Netherlands, Austria and Germany. Sampling points were evenly distributed in the Netherlands. In Germany, the majority of the samples were collected in Eastern Germany, Bavaria and North Rhine-Westphalia. One sample was located in Austria, close to the German border (Figure 1A).

The anaerobic digesters analyzed varied in terms of reactor type and general configuration. As shown in Figure 1B, over 50% of all samples were taken from continuously stirred tank reactors (CSTRs). This high number of CSTR samples is explained by the fact that it is the most common reactor type in the biogas sector (European Biogas Association, 2021). Although most plants are CSTRs, they differ in terms of reactor configurations, size, geometry, and agitation. Some used agitators, some used sprinklers, and some even used a floating roof, which, according to (Weiland, 2010), has the potential to stabilize the underlying microbiome. 13% of the samples were taken from plug flow reactors (PFRs). The PFR is designed to allow the feed to flow predominantly axial with minimal radial mixing, allowing for optimum residence time and contact between the feed and the microorganisms (Nasir et al., 2012). 10% of the samples were from industrial wastewater and wastewater treatment plants, and most of which used an upflow anaerobic sludge blanket reactor (UASB), which is commonly used for wastewater treatment (Lettinga et al., 1980). 6% of the samples were taken from two-staged AD systems (TSAD), in which the hydrolysis and the acidogenesis phases run separately from the acetogenesis and methanogenesis phases to meet the different requirements of the microorganisms (Holl et al., 2022). In order to cover a wider range of digester systems, some rarely occurring digester systems were sampled as well. These include leach bed systems, a unique peppercorn system and samples from other parts of the reactor system such as the secondary fermenter and feedstock. The samples used in this study are representative of the most common reactor systems in the biogas sector (Nasir et al., 2012).

The anaerobic digesters analyzed differed in terms of the collected operational process parameters (Supplementary Table 1 & 2). In addition to the reactor system, Figure 1 C shows four important operational parameters including temperature, hydraulic retention time, reactor volume and main substrate. The process temperature of the samples taken ranged between 25 °C and 65 °C, but the majority of the samples were between 35 °C and 50 °C, at the transition from mesophilic to thermophilic temperature. This range is advantageous for the process and is in line with other studies on the temperature of biogas plants (Hassa et al., 2021). Higher process temperatures lead to increased microbial activity and thus more biomass is converted to biogas per unit of time, which is why temperatures above 35 °C are beneficial (Pap et al., 2015). However, increases degradation rates that result from higher temperatures can lead to accelerated release of organic acids and other potentially process-inhibiting metabolites (Theuerl et al., 2018). The HRTs ranged from one days for the UASB and first stage of the TSAD systems to 170 days for the conventional CSTRs and secondary digesters. Although the range of HRTs is relatively broad, most reactors have an HRT between 15 and 60 days. Interestingly, small reactors tended to have low HRT, while substrates also had an impact on HRTs. In this regard, Figure 1 C shows that organic waste, sewage sludge, and industrial wastewater had short HRTs of about 20 days, which is consistent with the literature and suggests that these substrates can be converted quickly (Tufaner & Avşar, 2016). This is followed by a range between 20 and 80 days for energy crops, which tend to have higher fiber contents and therefore take longer to degrade (Kainthola et al., 2019). The last group are the residual crops including grass silage, wheat straw and green waste, which required HRTs of 80-180 days due to the high proportion of lignin and fiber components that are difficult to degrade (Appels et al., 2011). The use of manure or sewage sludge as a part of the substrate was found in all digesters as it is a well-suited seed sludge, which is already enriched in the key organisms for the methanogenic biogas process (Wu et al., 2010). The AD systems investigated in this study show a high diversity in terms of substrates used, reactor systems and operational parameters (Supplementary Table 1). This diversity sets this study apart from previous studies (Hassa et al., 2018; Kirkegaard et al., 2017; Theuerl et al., 2020). In addition to the main substrates mentioned above, rare cases with substrates such as slaughterhouse waste, grease, industrial wastewater, supermarket waste, unique residual materials such as sudan grass, sugar beet, millet and manures from goats and horses, were used (Supplementary Table 1). All the mentioned parameters such as reactor systems and configurations, temperature, retention time, volume, and feedstock underline the high diversity of the here presented set of samples. Samples from this study represent the majority of industrially applied plant types in the biogas sector. In combination with the respective chemical and taxonomic data, one of the most holistic data sets on biogas plants up to now is provided. Overall, a comprehensive range of chemical and operational parameters of these digesters were determined in accordance with the literature (Abendroth et al., 2015; Hao et al., 2016; Vrieze et al., 2016) and are shown in the Supplementary Table 1 and 2.

### 3.2 Chemical parameters of the anaerobic digestion systems

A comprehensive set of chemical parameters was collected for all the samples analyzed in this study (Figure 2A, Supplementary Table 2, 3). Due to the wide variety of operational process parameters, high variations of the measured chemical parameters were observed. Total volatile fatty acids (TVFAs) exhibited a large upward scatter, which could be explained due to the different systems, their specific configurations, varying feeding conditions and individual conditions in respect to the physical-chemical parameters (Figure 2A). The strong variations of key parameters such as VFAs among reactors indicate that the sampled reactors bear a high diversity of microbial habitats. Generally, one-stage systems are operated and designed to keep VFAs as low as possible. In contrast, the first stage of TSAD reactors have high concentrations of VFAs. In such systems, it is aimed to separate hydrolysis/acidogenesis from acetogenesis/methanogenesis to optimize the respective process conditions. To suppress hydrolysis/acidogenesis, the HRT is maximized. As a result, VFA concentrations increase and the pH drops, which impairs methanogenic archaea and related acetogens. The VFAs produced from the first stage are then fed into a second stage, the methane stage. This concept has been investigated already for 50 years (Holl et al., 2022) and, interestingly, Holl et al. mention that 50 years of research have not led to industrial solutions. This is not quite correct, since three industrial TSAD systems were investigated in the present study. However, chemical parameters (Supplementary Table 2) revealed the presence of methanogenesis in the first stage despite high loading rates. For example, and to the surprise of the respective plant operators, the pH was above 7.0 in all hydrolysis/acidogenesis stages, which is optimal for methanogenesis. One of the plant operators highlighted that they had noticed high ratios of methane in the hydrolysis/acidification stage, which are not expected there. The operator tried to prevent this by increasing the organic loading rate further, however with a maximal OLR of 44.6 kg⸱m^-3^⸱d^-1^, there were still methanogens which indicates, that they were able to adapt due to adaptive evolution. A further increase in the OLR was not possible without overloading the pumps of the respective plants. Thus, the statement by Holl et al. could be reformulated in such a way that although several two-stage industrial-scale plants in have already been built, no industrial solutions are known which clearly follow the concept of a two-stage biogas plant in terms of process technology. Apart from TSADs, high VFA concentrations were also observed in plug flow reactors. Although these reactors are not considered as classical two-stage systems, they still have high levels of VFAs due to the high OLR. For this reason, all plug flow reactors sampled in this study have a downstream methane stage to increase the degradation efficiency.

**Figure 2:**
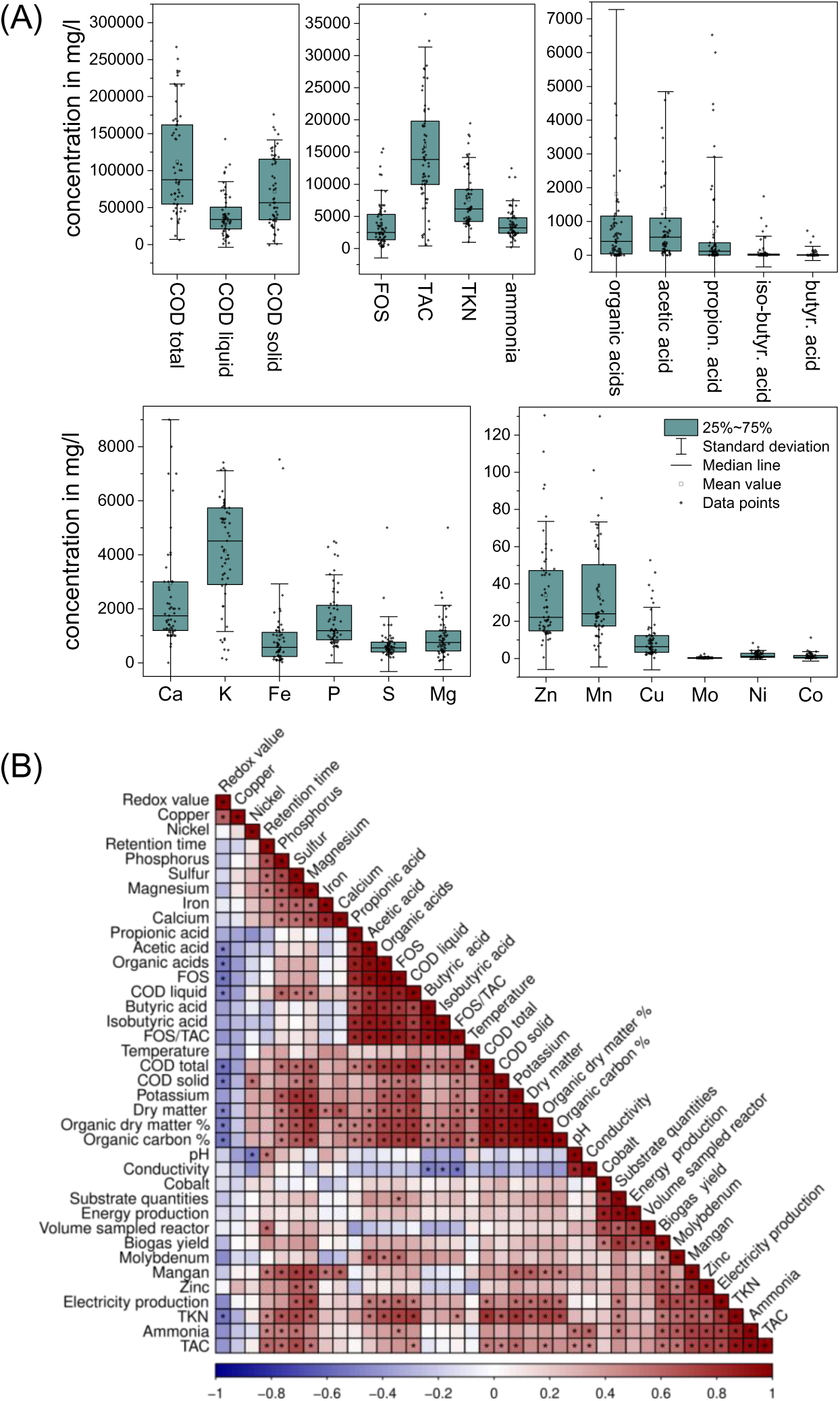
(A) Boxplots depicting the concentration of the main chemical parameters. Box size indicates the distribution of values between 25 and 75% of the values. (B) Spearman correlation analysis comparing the chemical variables. Positive correlations (i.e., positive Spearman’s rank correlation coefficients) are highlighted in red, negative correlations are highlighted in blue and the black dots indicate whether the respective correlation is significant.

Despite the large upward scatter in respect to VFAs, most digesters had a rather low concentration of VFAs. Therefore, it can be assumed that there is no significant inhibition of the methanogenic activity due to VFAs. The average acetate concentration was 1373 mg/l, which is far below inhibiting conditions of ∼2400 mg/l for one stage systems (Franke-Whittle et al., 2014). Despite maximum OLR and high VFA concentrations in two-stage systems, long-term inhibition of methanogenesis was not observed there. The threshold mentioned by Franke-Whittle et al. is therefore dependent on the type of plant and on the degree of adaptation of methanogenic archaea. The average butyrate concentration was 52 mg/l, well below inhibiting conditions, which start at 1800 mg/l. Propionic acid was the VFA that was closest to an inhibiting range, with an average concentration of 716 mg/l (inhibition usually starts at 900 mg/l) (Franke-Whittle et al., 2014; Y. Wang et al., 2009). All other parameters showed lower deviations and thus a lower dispersion of the distributions of the 25-75% quantile, but with a tendency towards higher values. It must be emphasized that the values at which inhibition occurs vary depending on many factors, for example due to microbial adaptations and the degree of protonation of the VFA salts (Bischofsberger et al., 2005).

A wide dispersion can be also observed for the COD, the COD from the solid’s fractions, as well as for the total inorganic carbon (TAC), Ca, K and P. According to the literature, these parameters are strongly influenced by the OLR, i.e. mainly by the substrate, the substrate quantities and the hydraulic retention time (Collivignarelli et al., 2017; Sánchez et al., 2004). In contrast, the quantile for COD liquid, volatile fatty acids determined by two-point titration (FOS), total nitrogen according to Kjeldahl (TKN), ammonia and all other trace elements is narrow. Strong deviations from the norm of these parameter values are often accompanied by process disturbances. Therefore, these parameters are intended to be kept constantly low (Ajayi-Banji & Rahman, 2022; Bischofsberger et al., 2005). Furthermore, most of them are limited by physicochemical effects such as solubility limits, precipitation and pH dependence. In addition to the parameters shown in Figure 2, the following additional parameters were determined and presented as the mean with standard deviation of all samples: pH = 7.6 ± 0.4; ORP = -374 mV ± 77; conductivity = 16.6 µS/cm ± 8.5; total solids (TS) = 10.1% ± 6.7; and total organic solids (oTS) = 6.5% ± 3 (Supplementary Table 2). Compared to the literature, it is noticeable that the values for total COD and COD solids are comparatively high and more variable (Świątczak et al., 2019). The parameters TS and oTS also show high values, which can be explained by the utilization of plant-based material like energy crops and residual crops in the agricultural biogas plants (Modenbach & Nokes, 2012). The correlation between high TS and high COD has been demonstrated using a Spearman correlation analysis (Figure 2B) and is consistent with the literature (Nagao et al., 2012). It is striking that there are predominantly significant positive correlations among the parameters (Figure 2B), which can be explained by the cascade nature of the process in which different processes build on each other (H. Wang et al., 2016). These positive correlations include dependencies between the individual VFAs, which can be explained by the fact that VFAs are degraded stepwise from medium-length fatty acids to the short-chain organic acids. (Amani et al., 2011; Gabris et al., 2015; Jha & Schmidt, 2017). In addition, acids correlate positively with COD, TS and oTS and TKN. All of these parameters are indicators of nutrient availability, so it is reasonable to assume that an increase in nutrient availability will also lead to an increase in hydrolytic activity and the formation of more VFAs. All values that directly or indirectly indicate the amount of organic matter, such as COD, TS, oTS, correlate positively with TKN and the trace elements P, S, Mg and K. Both trace substances and nitrogen naturally occur in the substrate, especially in manure and plant components, thus explaining this correlation. In particular, high levels of these elements can be detected in the solid components. The trace elements Ni, Mn and Mo also showed significant positive correlations with each other. As many plant operators use additives containing these trace elements, this correlation could be expected. Negative correlations between chemical variables mainly concerned the site parameters, i.e. pH, conductivity and redox value, with only the redox value showing significant negative correlations. pH and conductivity showed little or no significant correlations with all parameters. This could be because these parameters can be influenced by many other factors i.e. pH is influenced by all acids, bases and buffer systems present in the reactor.

Overall, the chemical data confirm that the samples constitute a very heterogeneous data set. In particular, parameters such as organic acids, COD and some trace elements showed a wide range of possible values, displaying many significant positive correlations according to the Spearman correlation analysis, but without exceeding the inhibition values indicated in the literature.

### 3.3 Taxonomic profiling of the AD systems

A total of 42,939 different amplicon sequence variants (ASVs) were detected in the entire dataset, accounting for 1,858 genera and 61 phyla. The average richness and Shannon indices at the ASV level were 1,019.13 ± 279.65 and 4.43 ± 0.76, respectively. At the phylum level, the average AD reactor was dominated by *Bacillota* (53.5%) followed by *Bacteroidota* (10.0%) and the archaeal phylum *Euryarchaeota* (10.1%). *Actinomycetota* (6.4%), *Pseudomonadota* (previously *Proteobacteria*) (5.2%), *Synergistota* (5.3%) and *Chloroflexi* (3.0%) were also detected in lower abundances (Figure 3C). Thus, in our samples 4 phyla covered 80% of the population and 6 phyla covered 90% of the population. These results are consistent with previous findings reporting that *Bacillota*, especially *Clostridia* and *Bacilli* classes, and *Bacteroidota* dominate AD processes (Basak et al., 2022; Schnürer, 2016). *Pseudomonadota* and *Synergistetes* phyla are usually found in lower abundances but are still prevalent (Schnürer, 2016), as well as *Acidobacteria*, *Actinomycetota* or *Chloroflexi*.

**Figure 3:**
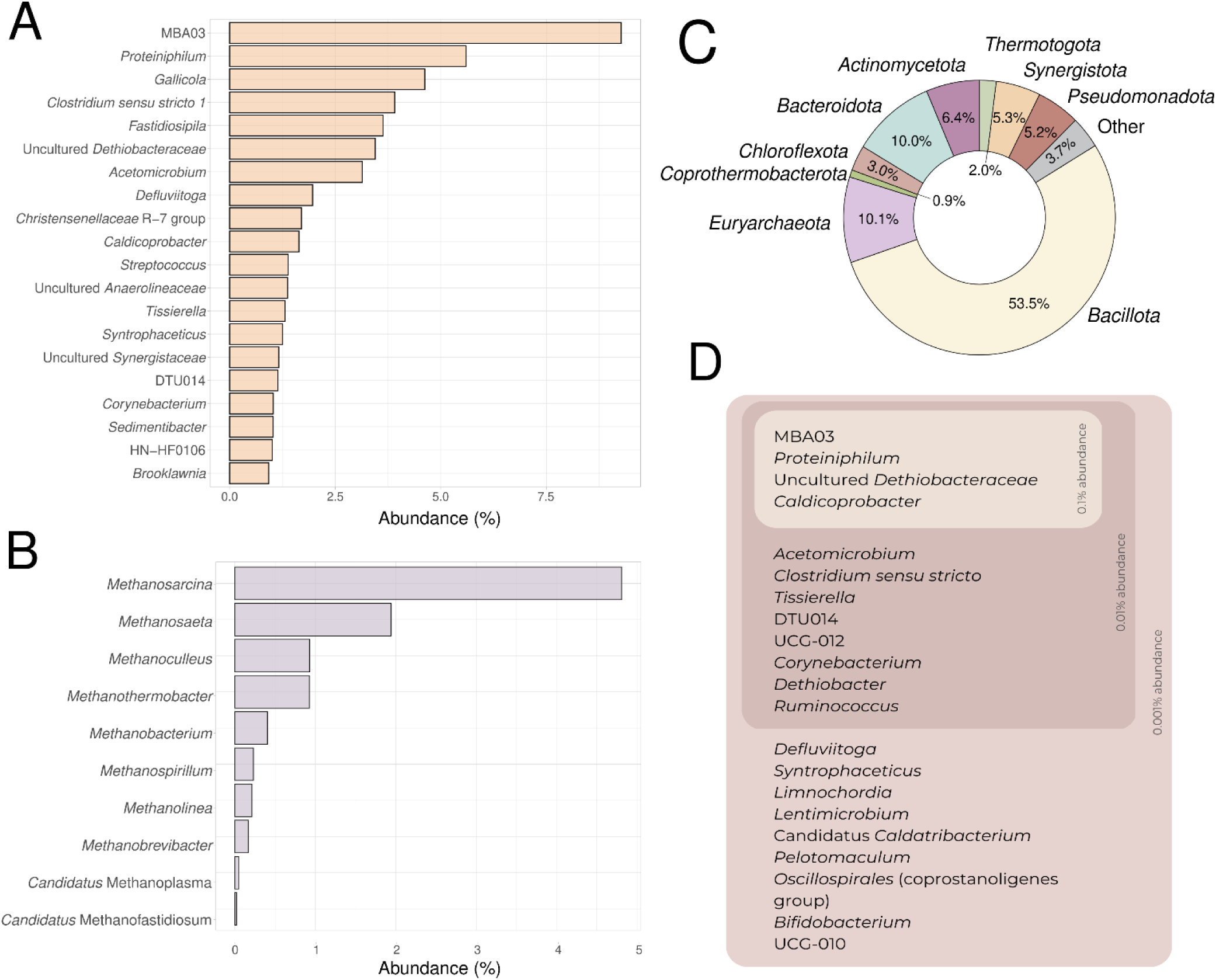
Mean abundances of the most abundant genera of: (A) Bacteria; (B) Archaea; (C) 10 most abundant phyla within the samples; and (D) core microbiome of the 80 samples analyzed, at 99% prevalence and different abundances: 0.001% (outer layer), 0.01% (mid layer) and 0.1% (inner layer).

At the genus level, taxonomic profiles were dominated by MBA03 (9.3%), followed by *Proteiniphilum* (5.6%), *Gallicola* (4.6%) and *Clostridium* sensu stricto (3.9%). Other abundant genera were *Acetomicrobium*, an uncultured *Dethiobacteraceae*, *Syntrophaceticus*, DTU014 and *Caldicoprobacter*, which were also present in the core microbiome (Figure 3D, Supplementary Table 4A). *Archaea* represented nearly 10% of the total microorganisms in the samples, with a maximum abundance of 48% in one sample. *Methanosarcina* was on average the most abundant archaeal genera (4.8%). This genus was detected in 78/80 samples, indicating a high prevalence in full-scale reactors. Other abundant archaea were *Methanothrix* (1.9%), *Methanoculleus* (0.9%) and *Methanothermobacter* (0.9%) (Supplementary Table 4A). It is important to highlight that some archaeal genera were assigned to phylum *Halobacterota* according to the SILVA database (v. 138) (Quast et al., 2013), while the “List of Prokaryotic names with Standing in Nomenclature” (LPSN) (Parte et al., 2020) classifies them as members of the phylum *Euryarchaeota*. In these cases, the taxonomy was manually corrected to match de LPSN criteria.

Previous studies have also identified members of the orders *Methanosarcinales*, *Methanobacteriales* and *Methanomicrobiales* as the dominating methanogenic archaea in AD systems (Campanaro et al., 2016; Schnürer, 2016) Moreover, *Methanosarcinales* occurs in the core microbiome and can be detected in averages up to 5% (Murillo-Roos et al., 2022). According to the results, the MBA03 genera and *Caldicoprobacter* from the *Bacillota* phylum, the genus *Proteiniphilum* from *Bacteroidota* phylum, and an uncultured organism from *Dethiobacteraceae* family, compose the core microbiome of anaerobic digesters, as they are present in 100% of the processed samples with an abundance higher than 0.1% (Figure 3D). When looking at the functions of the detected core bacteria in AD, two main roles can be identified. On one hand, the microorganisms of the MBA03 genera, the DTU14 genera and a genus of the *Dethiobacteraceae* family show high ammonia tolerance and syntrophic acetate oxidation (SAO) activities, thus contributing to hydrogenotrophic methanogenesis (Perman et al., 2022). On the other hand, MBA03, *Caldicoprobacter*, *Proteiniphilum*, *Acetomicrobium* and *Defluviitoga* show mainly hydrolytic activities (Kim et al., 2018; Perman et al., 2022). MBA03, and *Defluviitoga* can degrade complex carbohydrates such as xylan, cellulose and lignocellulose (Jensen et al., 2021; S. Zhang et al., 2022). *Proteinipihlum* and *Acetomicrobium* can degrade both peptides and complex carbohydrates (Perman et al., 2022; Tomazetto et al., 2018; Z. Wu et al., 2021), and *Caldicoprobacter* is considered to have the ability to hydrolyze lipids, peptides and carbohydrates (Lei et al., 2019). It must be highlighted that MBA03 is the only taxon that performs both major functions in the system, displaying both syntrophic acetate oxidation (SAO) and cellulolytic and xylanolytic activities, potentially playing a key role in AD systems. (Jensen et al., 2021; Perman et al., 2022, Puchol et al. unpublished).

### 3.4 Multivariate Spearman correlation analysis

In this study, Spearman correlation analysis showed many associations between microorganisms and the analyzed chemical variables, whereas for the operational parameters, only temperature and substrate amounts showed significant correlations with the taxonomic profiles (Figure 4, Supplementary Table 4B). Hierarchical clustering revealed two clearly distinct clusters of microorganisms showing similar trends with respect to chemical variables. The first cluster showed a higher number of bacteria belonging to phyla *Pseudomonadota*, *Actinomycetota*, *Chloroflexi* and *Bacteroidota*, reporting negative correlations with all the organic acids, nitrogen and COD. These bacterial genera are involved in the hydrolysis by degrading complex molecules such as chitin, peptide, lignocellulose and/or cellulose (Koeck et al., 2014; Perman et al., 2022; Serna-García et al., 2020). When acidogenesis begins after the hydrolytic reactions, the concentration of organic acid begins to increase, while the concentration of complex macromolecules decreases. Consequently, the abundance of these bacteria may decrease accordingly.

**Figure 4:**
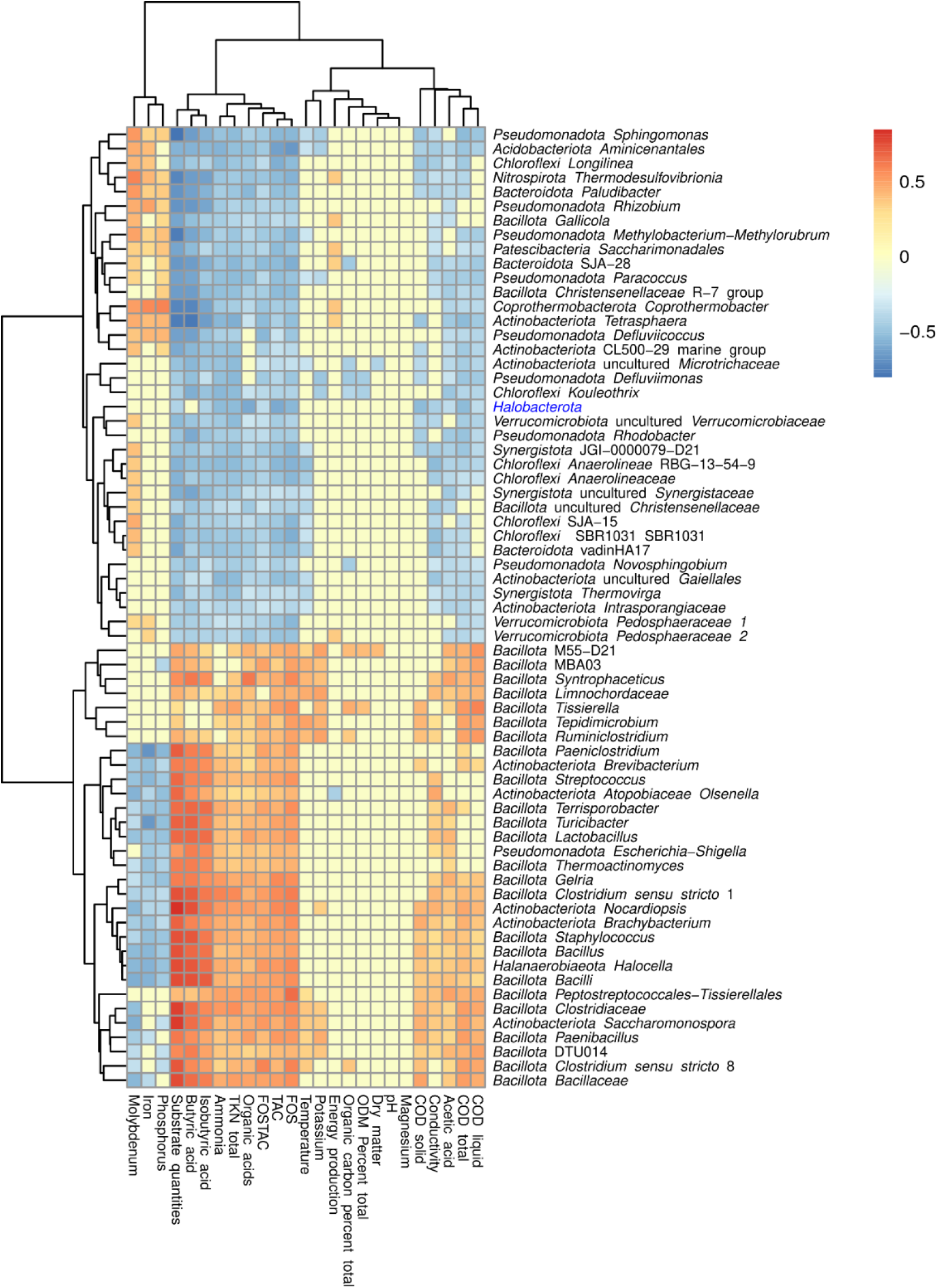
Spearman correlation analysis of 66 genera showing at least 14 correlations with chemical variables were plotted in a heatmap using the complete linkage method for hierarchical clustering. In this heatmap, red colors represent positive correlations while blue colors represent negative Spearman correlations. The full description of all the correlations detected are shown in Supplementary Table 4B. All the correlations plotted are significant (p-value <0.05). Halobacterota is highlighted because the Silva database incorrectly assigns the methanogen Methanoculleus to the genus. Archaea are highlighted in blue.

Both *Chloroflexi* and *Bacteroidota* are known to be typical acidogens with hydrolytic activity, capable of metabolising simple compounds such as amino acids, glycerol, glucose and complex polysaccharides (Azman et al. 2015; Chen et al. 2019; Perman et al. 2022). Within *Bacteroidota*, a higher presence of the *Paludibacteraceae* family (i.e. *Paludibacter* genera) has been reported in the earlier steps of AD processes (Perman et al., 2018)(Perman et al., 2022)^OBJOBJOBJ^. Members of the *Anaerolineaceae* family (i.e., *Longilinea*, *Anaerolinea*), which is the most abundant in the *Chloroflexales* order, metabolise polysaccharides such as pectin and xylan, and produce acetic and lactic acid and hydrogen (Zhang et al., 2018) (L. Zhang et al., 2018)^OBJOBJOBJOBJ^

Some microorganisms belonging to the *Bacillota* phylum, such as *Gallicola,* were also found in the first cluster, negatively correlating with organic acids. These non-saccharolytic genera can metabolize peptone and amino acids to organic acids (Cardinali-Rezende et al., 2016; Perman et al., 2022; Shi et al., 2021). There may be two reasons for this negative correlation: first, *Gallicola* can occur in a small window of organic acid concentration, and with increased organic acid content, the activity decreases; and second, *Gallicola* only metabolizes peptides, so its occurrence is limited by the presence of peptides (Cardinali-Rezende et al., 2016).

The second cluster of microorganisms showed positive correlations with organic acids (i.e., acetic acid, butyric acid and iso-butyric acid), ammonia, nitrogen, COD, substrate quantities and negative correlations with Fe, P and Mo. According to some studies, the *Syntrophaceticus* genera, MBA03, DTU014 and the *Dethiobacteraceae* family within the *Bacillota* phylum are potential syntrophic acetate oxidizing bacteria (SAOB) (Li et al., 2022; D. Zheng et al., 2019; Z. Zheng et al., 2021). Therefore, these bacteria would oxidize acetate to CO_2_ and H_2_, which would then be used to produce methane by hydrogenotrophic archaea. In addition, the orders DTU014 and MBA03 and members of the *Dethiobacteraceae* family have been reported to increase their abundance with the progression of AD (Perman et al., 2022), which fits with the hypothesis that the genera in the first cluster represent microorganisms performing hydrolysis and acidogenesis, while the microorganism in the second cluster are involved in the steps closer to acetogenesis and methanogenesis.

Correlation analysis also showed which prokaryotes were influenced by the abundance of other microorganisms, providing relevant information about the relationship between taxa during the AD processes (Fig 5, Supplementary Table 4C). The *Methanothermobacter* genera, *Methanoculleus* genera and *Methanosarcina* genera, which are hydrogenotrophic methanogens (Schnürer, 2016), positively correlated with potential SAOB, such as DTU014, MBA03, *Syntrophaceticus* and *Dethiobacter*, forming a cluster (Figure 5), whereas *Methanothrix,* an acetoclastic methanogen, negatively correlated with these bacterial genera. This supports the hypothesis that the abundance of SAOB is tightly related with hydrogenotrophic archaea; these bacteria would compete for acetate with acetoclastic methanogens and use acetate to generate hydrogen, thus producing a shift of the balance towards hydrogenotrophic methanogenesis (Schnürer, 2016; Westerholm et al., 2016) Most *Chloroflexota* genera, mainly dominated by the *Anaerolineaceae* family, negatively correlated with the genera in the cluster of syntrophs (DTU014, MBA03, *Syntrophaceticus*, *Clostridium sensu stricto* 1), while displaying positive correlations with *Christensenellaceae*-R7-group and *Gallicola*. Some bacteria within the syntrophic cluster were also positively correlated with temperature (Figure 4), indicating that their growth is favored in thermophilic conditions. In this sense, members of class *Clostridia* are known to be more abundant in high temperatures and increasing ammonia levels (Vrieze et al., 2015). Although high levels of ammonia can cause the inhibition of the process, SAOB seemed to be particularly tolerant to ammonia (Figure 4), and this may result in a shift to hydrogenotrophic methanogenesis, since these methanogens grow in syntrophy with SAOs (Schnürer, 2016). Interestingly, other authors have also reported positive correlations between ammonia and hydrogenotrophic methanogens and SAOB (Vrieze et al., 2015; Westerholm et al., 2016).

**Figure 5:**
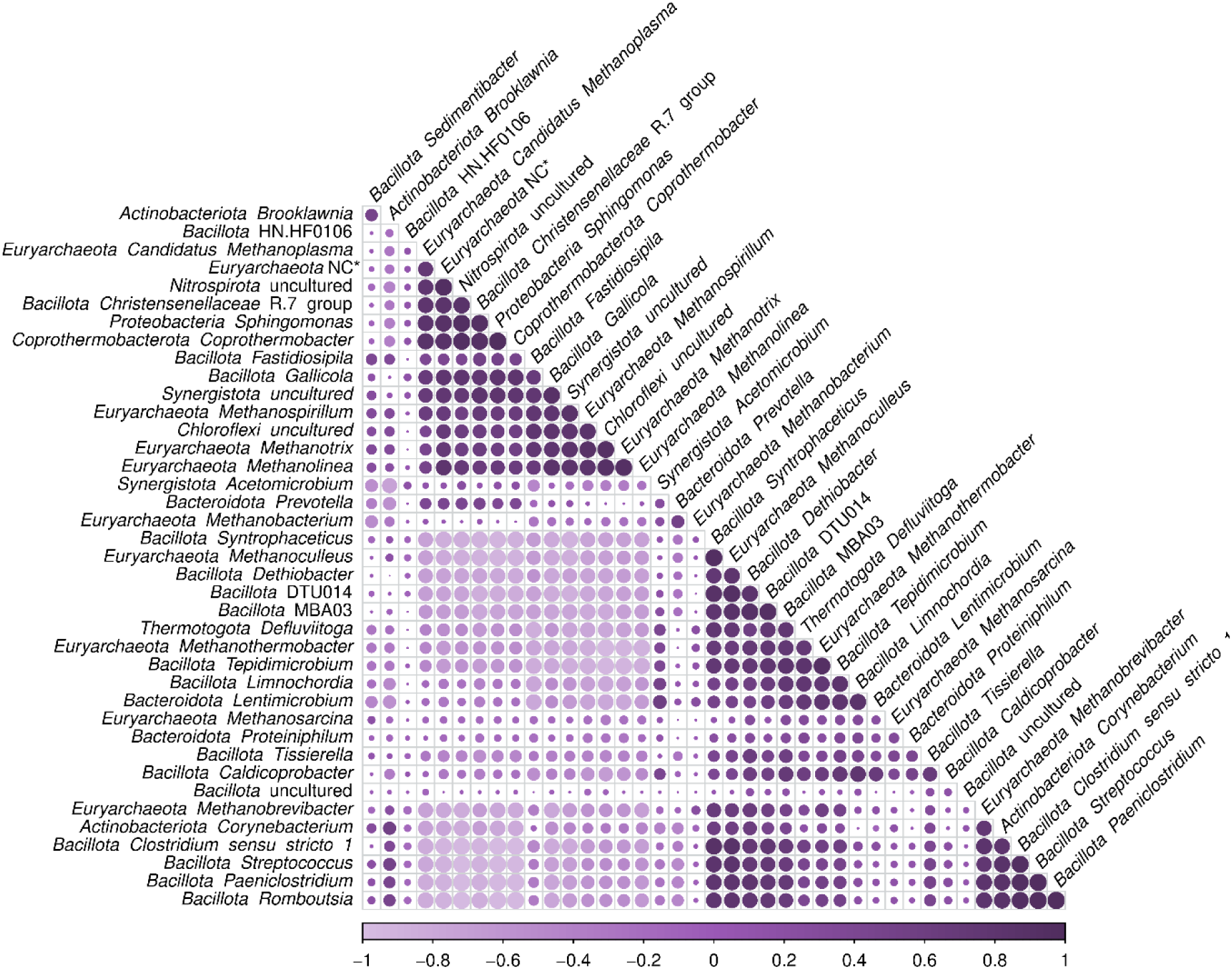
Correlations between the most abundant 30 bacteria and 10 archaea. The different shades of purple show the value of the correlation (legend below), while the size of the dot shows the significance of the correlation. The full description of all the correlations detected are shown in Supplementary Table 4C.

### 3.5 Influence of temperature and nitrogen content on the microbiome

Both temperature and ammonia variables were used as categorical variables for further comparisons. As reported by several authors, a reactor was considered “mesophilic” when displaying operational temperatures up to 45 °C, and “thermophilic” if the temperature was above 45 °C (Schnürer, 2016; Yenigün & Demirel, 2013).

In total, 41 genera were significantly more abundant in thermophilic conditions, while 171 increased their abundance in mesophilic conditions (Supplementary Table 4D). Different archaea, such as *Methanomethylovorans*, *Methanothrix* (previously *Methanosaeta*), *Methanolinea* and *Methanospirillum,* were more abundant in mesophilic conditions. The methanogenic community is especially sensitive to process instability (Madigou et al., 2019; Schnürer, 2016), and thermophilic reactors have been reported previously to be less stable in comparison to mesophilic ones (Lee et al., 2017). Moreover, the *Fastidiosipila* genera and the *Petrimonas* genera were more abundant in mesophilic conditions, in accordance with previous reports (Kim et al., 2018).

According to the results obtained (Figure 4), the most abundant microorganisms in thermophilic conditions belong to the *Bacillota* phylum (70% of them to class *Clostridia*). Specifically, thermophilic reactors were enriched in MBA03, DTU014, *Syntrophaceticus*, *Lentimicrobium*, *Defluviitoga* and *Tepidimicrobium* (Figure 6). The latter two bacteria have been found to be overexpressed in thermophilic reactors in previous studies (Kim et al., 2018).

**Figure 6:**
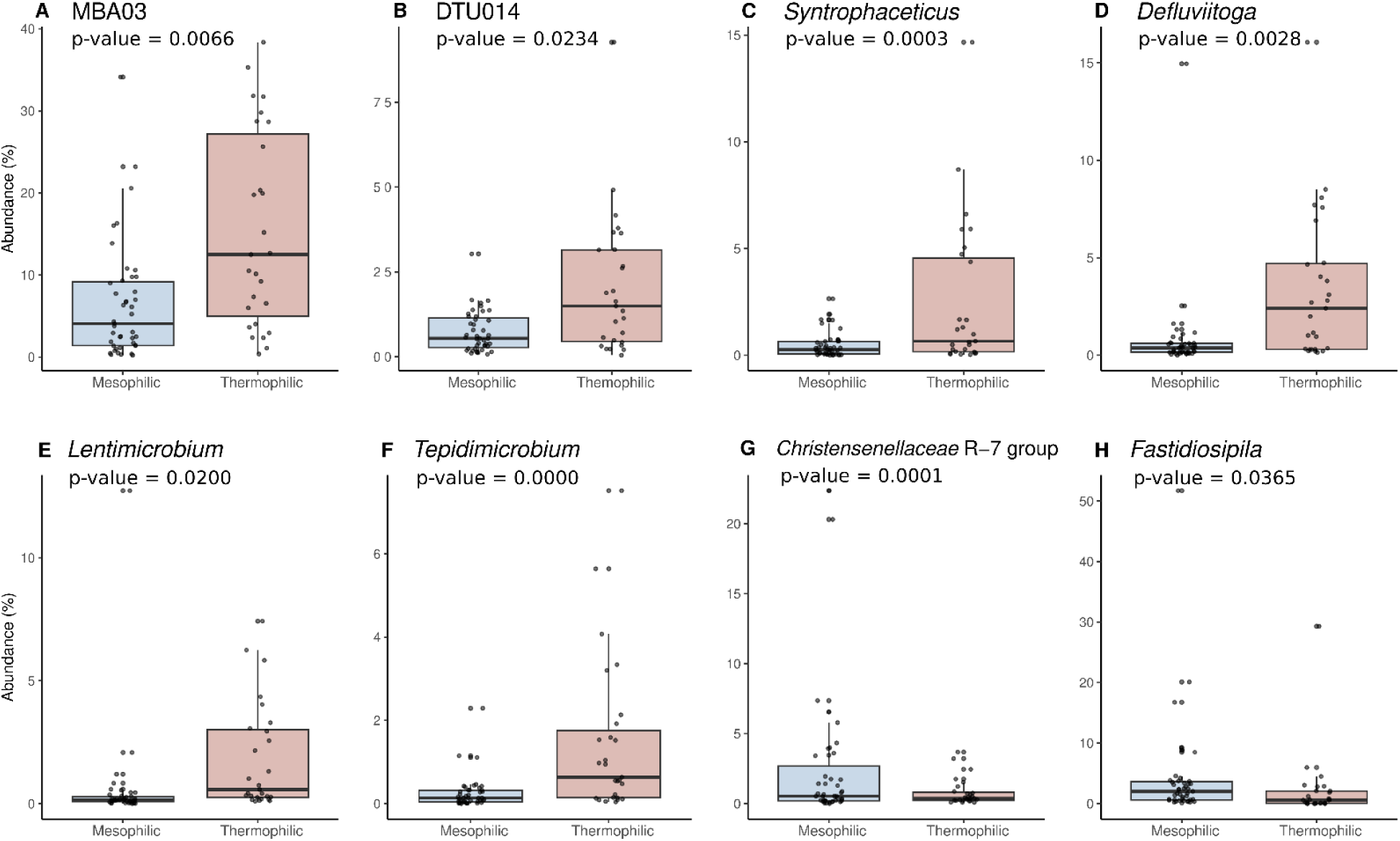
Differential expression of eight bacterial genera in different temperature conditions. Blue represents mesophilic condition (<45°C) and pink represents thermophilic condition (≥45°C). The statistical analysis was performed with DESeq2 in RStudio.

It is well known that high amounts of nitrogen, especially ammonia, can inhibit the AD process (Jiang et al., 2019; Yu et al., 2020). However, there is no consensus about the threshold of concentration of this molecule that starts negatively affecting biogas production, as the ammonia concentration depends on the pH, temperature and the processing state of the ammonia, which also depends on the substrate and the underlying microbiome. Some authors have previously reported that a critical threshold concentration of ammonia that causes toxicity or inhibition is between 1500 and 7000 mg/l (Jiang et al., 2019). To prove the effect of ammonia levels on the microbial community, the quantitative variable was divided into “low ammonia content” for values up to 5000 mg/l and “high ammonia content” for values above 5000 mg/l and the differential abundance in both situations was calculated (Supplementary Table 4E).

In low ammonia conditions, a higher abundance of archaeal genera (such as *Methanosarcina*, *Methanothrix*, *Methanospirillum*, *Methanoplasma* or *Methanolinea*) was detected. There is previous evidence that high ammonia conditions can affect the whole microbial community, but methanogenic archaea are the ones which suffer this stress the most, particularly acetoclastic methanogens (Feng et al., 2022; Westerholm et al., 2016).

Regarding the bacterial community, *Clostridium sensu stricto* 1, *Proteiniphilum* and *Defluviitoga*, among others, were more abundant in reactors with ammonia concentrations above 5000 mg/l (Figure 7). Both *Clostridium sensu stricto* 1 and *Proteiniphilum* have been reported to be abundant in high ammonia conditions (Feng et al., 2022; C. Wang et al., 2022) This could be due to their main ability to degrade proteins/amino acids (Perman et al., 2022). On the other hand, *Acetomicrobium*, *Christensenellaceae* R7 group and *Gallicola* were more abundant in lower ammonia conditions (Figure 7).

**Figure 7:**
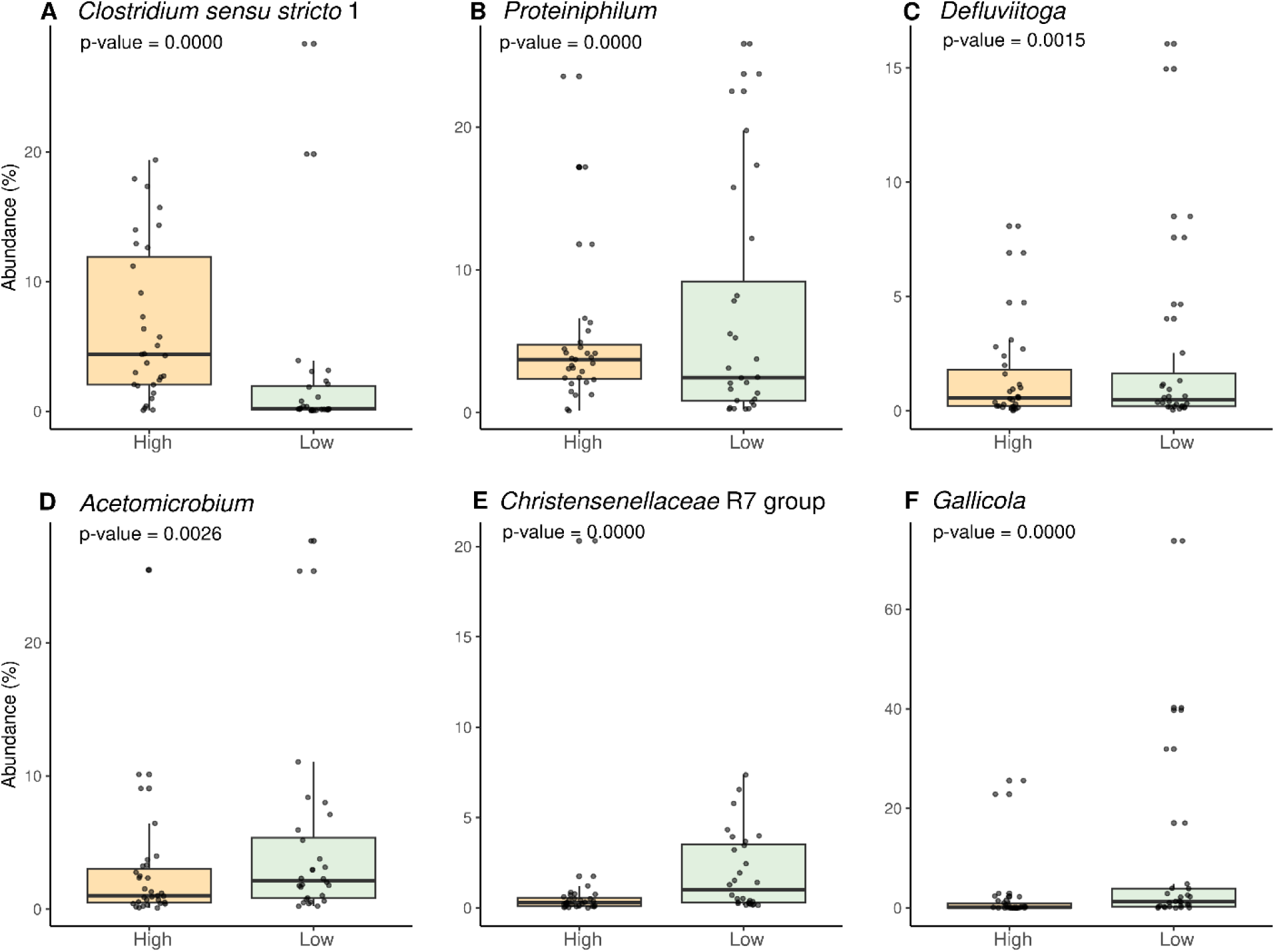
Differential expression of six bacterial genera in reactors with different nitrogen content. Yellow represents high ammonia content (>5000 mg/l) and green represents low ammonia content (<5000 mg/l). The statistical analysis was performed with DESeq2 in RStudio.

### 3.6 Relationship between microbial and technical variables

In order to study the relationship between the microbial communities and the substrates used in AD, the feedstocks were divided in three different groups, based on their chemical composition (Theuerl et al., 2018): organic biological waste (cluster “biowaste”), energy crops and agricultural animal waste (cluster “agricultural waste”) and wastewater sludge and industrial wastes (cluster “industrial waste”). It is important to highlight that the feedstocks used in full-scale reactors are a mixture of substrates from different sources, which complicated the analysis. However, some relevant conclusions were obtained.

First, the *Pseudomonadota* phylum was more abundant in the AD systems inoculated with substrates coming from industrial waste; this phylum has been previously related with UASB reactors treating sludges (Schnürer, 2016). Moreover, some core genera such as MBA03 and UCG-010, together with the hydrogenotrophic archaea *Methanoculleus* showed higher abundances in agricultural wastes and biowaste-based feedstock, while *Syntrophaceticus*, *Clostridium sensu stricto* 1 and the hydrogenotrophic archaea *Methanothermobacter* were more abundant in substrates from agricultural origins (Supplementary Table 4F) which was also reported in the literature (Niu et al., 2013).

Operational design of the AD process is another factor affecting the resulting microbial community and the final biogas yield. Different parameters, such as shear, granule formation, hydraulic retention time, liquid upflow velocity or feed rate (Kundu et al., 2017) can be determining for one microbial community to find their optimal conditions for growth.

Regarding reactor type, the most common ones among the samples were continuously stirred tank reactors (CSTR; 44 samples), plug flow fermenters (10 samples), upflow anaerobic sludge blanket digesters (UASBD; 8 samples) and two stage reactors with hydrolysis (5 samples). The differential expression analysis revealed that *Syntrophaceticus* and *Caldicoprobacter* were overexpressed in CSTRs, *Coprothermobacter* was overexpressed in plug flow fermenters, while other microorganisms present in the core microbiome, such as *Fastidiosipila*, *Gallicola* and *Christensenllaceae* R7 group were more abundant in UASB digesters (Supplementary Figure 4G).

Two hydrogenotrophic archaea, *Methanothermobacter* and *Methanolinea*, were overexpressed in CSTRs and two stage reactors, respectively, while *Methanothrix* was more abundant in UASB reactors. Genera from the *Pseudomonadota* phylum were particularly abundant in UASB, which is in line with the results obtained for substrates coming from industrial wastes and with previous developments (Y. Wu et al., 2016; H. Xu et al., 2020).

### 3.6 Limitations & outlook

Despite research efforts made in recent years, AD is still a microbial black box due to the complexity of the microbial transformations and interactions, the variability of process designs, and the high number of technical and chemical variables that affect the underlying microbiomes.

For this reason, the present study faced several challenges. First, the substrates used in full-scale reactors were diverse and complex, consisting mainly of mixtures of different feedstocks in unknown proportions, which made it difficult to determine the true influence of the substrate on the AD microbiomes (J. Zhang et al., 2017). Moreover, this study focused on studying bacteria and archaea, but viruses (J. Zhang et al., 2017), protozoa (Priya et al., 2008) or anaerobic fungi (Kazemi Shariat Panahi et al., 2022), are also important members of the AD microbiomes and their role in industrial biogas production should be addressed in future works. Finally, it must be highlighted that not all the industrial plant operators measured the amount of biogas produced in their systems. Also, the methods for the measurement of biogas and methane ratio differ, which make it difficult to compare such values between plants. This prevented the establishment of correlations between methane production/efficiency and the different chemical, taxonomic and technical parameters. This highlights the need to establish standardized methods for methane measurement that can be easily adopted by industrial plant operators, thus it would allow for the process to be controlled in a more robust way, and enable the possibility of evaluating the effect of bioaugmentation or biostimulation approaches on biogas production.

## 4 Conclusions

The core microbiome of the studied full-scale anaerobic digesters consisted of MBA03, *Proteiniphilum*, a member of *Dethiobacteraceae* and *Caldicoprobacter*. *Methanosarcina* was detected in 98% of samples. Several chemical and technical parameters like ammonium, temperature, COD, organic acids, reactor system, substrate selection and quantities affected the composition of the microbiomes. Spearman analyses revealed two clusters of microbes: the first included microorganisms that were associated with acetoclastic archaea, while the other was formed by hydrogenotrophic archaea and several potential SAOB. Overall, this study provides a diverse, holistic and comprehensive dataset on the operation of 45 full-scale anaerobic digesters and sheds light on the role of various chemical, technical and microbiological variables in the biogas production process.

## Acknowledgements

Funding information: Financial support from the European Union (MICRO4BIOGAS project with reference ID101000470 funded by European Union’s Horizon 2020 research and innovation programme) is acknowledged.

## Supplementary information

Supplementary data associated with this article can be found, in the online version.

· Supplementary Tables 1-3

· Supplementary Tables 4A-4G

· Supplementary Figure 1-3

## Notes

### Competing Interest Statement

The authors have declared no competing interest.

## References

Abendroth, C., Vilanova, C., Günther, T., Luschnig, O., & Porcar, M. (2015). Eubacteria and archaea communities in seven mesophile anaerobic digester plants in Germany. Biotechnology for Biofuels, 8, 87. 10.1186/s13068-015-0271-6

Ajayi-Banji, A., & Rahman, S. (2022). A review of process parameters influence in solid-state anaerobic digestion: Focus on performance stability thresholds. Renewable and Sustainable Energy Reviews, 167, 112756. 10.1016/j.rser.2022.112756

Amani, T., Nosrati, M., Mousavi, S. M., & Kermanshahi, R. K. (2011). Study of syntrophic anaerobic digestion of volatile fatty acids using enriched cultures at mesophilic conditions. International Journal of Environmental Science & Technology, 8(1), 83–96. 10.1007/BF03326198

Appels, L., Lauwers, J., Degrève, J., Helsen, L., Lievens, B., Willems, K., van Impe, J., & Dewil, R. (2011). Anaerobic digestion in global bio-energy production: Potential and research challenges. Renewable and Sustainable Energy Reviews, 15(9), 4295–4301. 10.1016/j.rser.2011.07.121

Azman, S., Khadem, A. F., van Lier, J. B., Zeeman, G., & Plugge, C. M. (2015). Presence and Role of Anaerobic Hydrolytic Microbes in Conversion of Lignocellulosic Biomass for Biogas Production. Critical Reviews in Environmental Science and Technology, 45(23), 2523–2564. 10.1080/10643389.2015.1053727

Basak, B., Patil, S. M., Kumar, R., Ahn, Y., Ha, G. S., Park, Y. K., Ali Khan, M., Jin Chung, W., Woong Chang, S., & Jeon, B. H. (2022). Syntrophic bacteria- and Methanosarcina-rich acclimatized microbiota with better carbohydrate metabolism enhances biomethanation of fractionated lignocellulosic biocomponents. Bioresource Technology, 360, 127602. 10.1016/j.biortech.2022.127602

Bischofsberger, W., Dichtl, N., Rosenwinkel, K. H., Seyfried, C. F., & Böhnke, B. (Eds.). (2005). Anaerobtechnik (2., vollständig überarbeitete Auflage). Springer. http://digitale-objekte.hbz-nrw.de/webclient/DeliveryMana-ger?pid=1501187&custom_att_2=simple_viewer

Bolyen, E., Rideout, J. R., Dillon, M. R., Bokulich, N. A., Abnet, C. C., Al-Ghalith, G. A., Ale-xander, H., Alm, E. J., Arumugam, M., Asnicar, F., Bai, Y., Bisanz, J. E., Bittinger, K., Brejnrod, A., Brislawn, C. J., Brown, C. T., Callahan, B. J., Caraballo-Rodríguez, A. M., Chase, J., … Caporaso, J. G. (2019). Reproducible, interactive, scalable and extensible microbiome data science using QIIME 2. Nature Biotechnology, 37(8), 852–857. 10.1038/s41587-019-0209-9

Calusinska, M., Goux, X., Fossépré, M., Muller, E. E. L., Wilmes, P., & Delfosse, P. (2018). A year of monitoring 20 mesophilic full-scale bioreactors reveals the existence of stable but different core microbiomes in bio-waste and wastewater anaerobic digestion systems. Bio-technology for Biofuels, 11, 196. 10.1186/s13068-018-1195-8

Campanaro, S., Treu, L., Kougias, P. G., Francisci, D. de, Valle, G., & Angelidaki, I. (2016). Metagenomic analysis and functional characterization of the biogas microbiome using high throughput shotgun sequencing and a novel binning strategy. Biotechnology for Bio-fuels, 9, 26. 10.1186/s13068-016-0441-1

Cardinali-Rezende, J., Rojas-Ojeda, P., Nascimento, A. M., & Sanz, J. L. (2016). Proteolytic bacterial dominance in a full-scale municipal solid waste anaerobic reactor assessed by 454 pyrosequencing technology. Chemosphere, 146, 519–525. 10.1016/j.chemosphere.2015.12.003

Chen, L., Liu, J [Junwei], Ge, X., Xu, W., Chen, Y., Li, F., Cheng, D., & Shao, R. (2019). Simulated digestion and fermentation in vitro by human gut microbiota of polysaccharides from Helicteres angustifolia L. International Journal of Biological Macromolecules, 141, 1065–1071. 10.1016/j.ijbiomac.2019.09.073

Collivignarelli, M. C., Bertanza, G., Abbà, A., Sordi, M., & Pedrazzani, R. (2017). Synergy between anaerobic digestion and a post-treatment based on Thermophilic Aerobic Membrane Reactor (TAMR). Environmental Progress & Sustainable Energy, 36(6), 1802– 1809. 10.1002/ep.12677

Emerson, K., Russo, R. C., Lund, R. E., & Thurston, R. V. (1975). Aqueous Ammonia Equilibrium Calculations: Effect of pH and Temperature. Journal of the Fisheries Research Board of Canada, 32(12), 2379–2383. 10.1139/f75-274

European Biogas Association. (2021). Statistical Report of the European Biogas Association 2021. Brussels.

Feng, G., Zeng, Y., Wang, H. Z., Chen, Y. T., & Tang, Y. Q. (2022). Proteiniphilum and Methanothrix harundinacea became dominant acetate utilizers in a methanogenic reactor operated under strong ammonia stress. Frontiers in Microbiology, 13, 1098814. 10.3389/fmicb.2022.1098814

Franke-Whittle, I. H., Walter, A., Ebner, C., & Insam, H. (2014). Investigation into the effect of high concentrations of volatile fatty acids in anaerobic digestion on methanogenic communities. *Waste Management (New York*, N.Y.), 34(11), 2080–2089. 10.1016/j.wasman.2014.07.020

Gabris, C., Bengelsdorf, F. R., & Dürre, P. (2015). Analysis of the key enzymes of butyric and acetic acid fermentation in biogas reactors. Microbial Biotechnology, 8(5), 865–873. 10.1111/1751-7915.12299

Golberg, A., Sack, M., Teissie, J., Pataro, G., Pliquett, U., Saulis, G., Stefan, T., Miklavcic, D., Vorobiev, E., & Frey, W. (2016). Energy-efficient biomass processing with pulsed electric fields for bioeconomy and sustainable development. Biotechnology for Biofuels, 9, 94. 10.1186/s13068-016-0508-z

Hansen, K. H., Irini Angelidaki, & and Birgitte Kiaer Ahring (1998). Anaerobic digestion of swine manure: inhibition by ammonia. Water Research, 32(1), 5–12. 10.1016/S0043-1354(97)00201-7

Hao, L., Bize, A., Conteau, D., Chapleur, O., Courtois, S., Kroff, P., Desmond-Le Quéméner, E., Bouchez, T., & Mazéas, L. (2016). New insights into the key microbial phylotypes of anaerobic sludge digesters under different operational conditions. Water Research, 102, 158–169. 10.1016/j.watres.2016.06.014

Hassa, J., Klang, J., Benndorf, D., Pohl, M., Hülsemann, B., Mächtig, T., Effenberger, M., Pühler, A., Schlüter, A., & Theuerl, S. (2021). Indicative Marker Microbiome Structures Deduced from the Taxonomic Inventory of 67 Full-Scale Anaerobic Digesters of 49 Agricultural Biogas Plants. Microorganisms, 9(7). 10.3390/microorga-nisms9071457.

Hassa, J., Maus, I., Off, S., Pühler, A., Scherer, P., Klocke, M., & Schlüter, A. (2018). Metagenome, metatranscriptome, and metaproteome approaches unraveled compositions and functional relationships of microbial communities residing in biogas plants. Applied Microbiology and Biotechnology, 102(12), 5045–5063. 10.1007/s00253-018-8976-7

Holl, E., Steinbrenner, J., Merkle, W., Krümpel, J., Lansing, S., Baier, U., Oechsner, H., & Lemmer, A. (2022). Two-stage anaerobic digestion: State of technology and perspective roles in future energy systems. Bioresource Technology, 360, 127633. 10.1016/j.biortech.2022.127633

Jensen, M. B., Jonge, N. de, Dolriis, M. D., Kragelund, C., Fischer, C. H., Eskesen, M. R., Noer, K., Møller, H. B., Ottosen, L. D. M., Nielsen, J. L., & Kofoed, M. V. W. (2021). Cellulolytic and Xylanolytic Microbial Communities Associated With Lignocellulose-Rich Wheat Straw Degradation in Anaerobic Digestion. Frontiers in Microbiology, 12, 645174. 10.3389/fmicb.2021.645174

Jetten, M. S., Stams, A. J., & Zehnder, A. J. (1992). Methanogenesis from acetate: a comparison of the acetate metabolism in Methanothrix soehngenii and Methanosarcina spp. FEMS Microbiology Letters, 88(3-4), 181–198. 10.1111/j.1574-6968.1992.tb04987.x

Jha, P., & Schmidt, S. (2017). Reappraisal of chemical interference in anaerobic digestion processes. Renewable and Sustainable Energy Reviews, 75, 954–971. 10.1016/j.rser.2016.11.076

Jiang, Y., McAdam, E., Zhang, Y [Yue], Heaven, S., Banks, C., & Longhurst, P. (2019). Ammonia inhibition and toxicity in anaerobic digestion: A critical review. Journal of Water Process Engineering, 32, 100899. 10.1016/j.jwpe.2019.100899

Kainthola, J., Kalamdhad, A. S., & Goud, V. V. (2019). A review on enhanced biogas production from anaerobic digestion of lignocellulosic biomass by different enhancement techniques. Process Biochemistry, 84, 81–90. 10.1016/j.procbio.2019.05.023

Kazemi Shariat Panahi, H., Dehhaghi, M., Guillemin, G. J., Gupta, V. K., Lam, S. S., Aghbashlo, M., & Tabatabaei, M. (2022). A comprehensive review on anaerobic fungi applications in biofuels production. The Science of the Total Environment, 829, 154521. 10.1016/j.scitotenv.2022.154521.

Kim, E., Lee, J [Joonyeob], Han, G., & Hwang, S. (2018). Comprehensive analysis of microbial communities in full-scale mesophilic and thermophilic anaerobic digesters treating food waste-recycling wastewater. Bioresource Technology, 259, 442–450. 10.1016/j.biortech.2018.03.079

Kirkegaard, R. H., McIlroy, S. J., Kristensen, J. M., Nierychlo, M., Karst, S. M., Dueholm, M. S., Albertsen, M., & Nielsen, P. H [Per H.] (2017). The impact of immigration on microbial community composition in full-scale anaerobic digesters. Scientific Reports, 7(1), 9343. 10.1038/s41598-017-09303-0

Koeck, D. E., Pechtl, A., Zverlov, V. V., & Schwarz, W. H. (2014). Genomics of cellulolytic bacteria. Current Opinion in Biotechnology, 29, 171–183. 10.1016/j.cop-bio.2014.07.002

Kundu, K., Sharma, S., & Sreekrishnan, T. R. (2017). Influence of Process Parameters on Anaerobic Digestion Microbiome in Bioenergy Production: Towards an Improved Understanding. BioEnergy Research, 10(1), 288–303. 10.1007/s12155-016-9789-0

Lahti, L., Shetty, S., & Salojarvi, J. (2017). Microbiome R package. Advance online publication. 10.18129/B9.bioc.microbiome

Lee, J [Joonyeob], Shin, S. G., Han, G., Koo, T., & Hwang, S. (2017). Bacteria and archaea communities in full-scale thermophilic and mesophilic anaerobic digesters treating food wastewater: Key process parameters and microbial indicators of process instability. Bioresource Technology, 245(Pt A), 689–697. 10.1016/j.biortech.2017.09.015

Lei, Z., Zhi, L., Jiang, H., Chen, R., Wang, X [Xiaochang], & Li, Y.Y. (2019). Characterization of microbial evolution in high-solids methanogenic co-digestion of canned coffee processing wastewater and waste activated sludge by an anaerobic membrane bioreactor. Journal of Cleaner Production, 232, 1442–1451. 10.1016/j.jcle-pro.2019.06.045

Leng, L., Yang, P., Singh, S., Zhuang, H., Xu, L., Chen, W. H., Dolfing, J., Li, D., Zhang, Y [Yan], Zeng, H., Chu, W., & Lee, P. H. (2018). A review on the bioenergetics of anaerobic microbial metabolism close to the thermodynamic limits and its implications for digestion applications. Bioresource Technology, 247, 1095–1106. 10.1016/j.biortech.2017.09.103

Lettinga, G., van Velsen, A. F. M., Hobma, S. W., Zeeuw, W. de, & Klapwijk, A. (1980). Use of the upflow sludge blanket (USB) reactor concept for biological wastewater treatment, especially for anaerobic treatment. Biotechnology and Bioengineering, 22(4), 699–734. 10.1002/bit.260220402

Li, C., He, P., Hao, L., Lü, F., Shao, L., & Zhang, H. (2022). Diverse acetate-oxidizing syntrophs contributing to biogas production from food waste in full-scale anaerobic digesters in China. Renewable Energy, 193, 240–250. 10.1016/j.renene.2022.04.143

Love, M. I., Huber, W., & Anders, S. (2014). Moderated estimation of fold change and dispersion for RNA-seq data with DESeq2. Genome Biology, 15(12), 550. 10.1186/s13059-014-0550-8

Madigou, C., Lê Cao, K. A., Bureau, C., Mazéas, L., Déjean, S., & Chapleur, O. (2019). Ecological consequences of abrupt temperature changes in anaerobic digesters. Chemical Engineering Journal, 361, 266–277. 10.1016/j.cej.2018.12.003

McBee, R. H. (1954). The characteristics of Clostridium thermocellum. Journal of Bacteriology, 67(4), 505–506. 10.1128/jb.67.4.505-506.1954

McMurdie, P. J., & Holmes, S. (2013). Phyloseq: An R package for reproducible interactive analysis and graphics of microbiome census data. PloS One, 8(4), e61217. 10.1371/journal.pone.0061217

Modenbach, A. A., & Nokes, S. E. (2012). The use of high-solids loadings in biomass pretreatment--a review. Biotechnology and Bioengineering, 109(6), 1430–1442. 10.1002/bit.24464

Murillo-Roos, M., Uribe-Lorío, L., Fuentes-Schweizer, P., Vidaurre-Barahona, D., Brenes-Guillén, L., Jiménez, I., Arguedas, T., Liao, W., & Uribe, L. (2022). Biogas Production and Microbial Communities of Mesophilic and Thermophilic Anaerobic Co-Digestion of Animal Manures and Food Wastes in Costa Rica. Energies, 15(9), 3252. 10.3390/en15093252

Nagao, N., Tajima, N., Kawai, M., Niwa, C., Kurosawa, N., Matsuyama, T., Yusoff, F. M., & Toda, T. (2012). Maximum organic loading rate for the single-stage wet anaerobic digestion of food waste. Bioresource Technology, 118, 210–218. 10.1016/j.biortech.2012.05.045

Nasir, I. M., Mohd Ghazi, T. I., & Omar, R. (2012). Anaerobic digestion technology in livestock manure treatment for biogas production: A review. Engineering in Life Sciences, 12(3), 258–269. 10.1002/elsc.201100150

Ni, J., Hatori, S., Wang, Y [Yin], Li, Y. Y., & Kubota, K. (2020). Uncovering Viable Microbiome in Anaerobic Sludge Digesters by Propidium Monoazide (PMA)-PCR. Microbial Ecology, 79(4), 925–932. 10.1007/s00248-019-01449-w

Niu, Q., Qiao, W., Qiang, H., & Li, Y. Y. (2013). Microbial community shifts and biogas conversion computation during steady, inhibited and recovered stages of thermophilic methane fermentation on chicken manure with a wide variation of ammonia. Bioresource Technology, 146, 223–233. 10.1016/j.biortech.2013.07.038

Pap, B., Györkei, Á., Boboescu, I. Z., Nagy, I. K., Bíró, T., Kondorosi, É., & Maróti, G. (2015). Temperature-dependent transformation of biogas-producing microbial communities points to the increased importance of hydrogenotrophic methanogenesis under thermophilic operation. Bioresource Technology, 177, 375–380. 10.1016/j.biortech.2014.11.021

Parte, A. C., Sardà Carbasse, J., Meier-Kolthoff, J. P., Reimer, L. C., & Göker, M. (2020). List of Prokaryotic names with Standing in Nomenclature (LPSN) moves to the DSMZ. International Journal of Systematic and Evolutionary Microbiology, 70(11), 5607–5612. 10.1099/ijsem.0.004332

Perman, E., Schnürer, A., Björn, A., & Moestedt, J. (2022). Serial anaerobic digestion improves protein degradation and biogas production from mixed food waste. Biomass and Bioenergy, 161, 106478. 10.1016/j.biombioe.2022.106478

Priya, M., Haridas, A., & Manilal, V. B. (2008). Anaerobic protozoa and their growth in biomethanation systems. Biodegradation, 19(2), 179–185. 10.1007/s10532-007-9124-8

Puig-Castellví, F., Cardona, L., Jouan-Rimbaud Bouveresse, D., Cordella, C. B. Y., Mazéas, L., Rutledge, D. N., & Chapleur, O. (2020). Assessment of the microbial interplay during anaerobic co-digestion of wastewater sludge using common components analysis. PloS One, 15(5), e0232324. 10.1371/journal.pone.0232324.

Quast, C., Pruesse, E., Yilmaz, P., Gerken, J., Schweer, T., Yarza, P., Peplies, J., & Glöck-ner, F. O. (2013). The SILVA ribosomal RNA gene database project: Improved data processing and web-based tools. Nucleic Acids Research, 41(Database issue), D590-6. 10.1093/nar/gks1219

Sánchez, E., Borja, R., Travieso, L., Colmenarejo, M. F., Chica, A., & Martín, A. (2004). Treatment of settled piggery waste by a down-flow anaerobic fixed bed reactor. Journal of Chemical Technology & Biotechnology, 79(8), 851–862. 10.1002/jctb.1059

Satari, L., Guillén, A., Vidal-Verdú, À., & Porcar, M. (2020). The wasted chewing gum bacteriome. Scientific Reports, 10(1), 16846. 10.1038/s41598-020-73913-4

Schnürer, A. (2016). Biogas Production: Microbiology and Technology. Advances in Biochemical Engineering/biotechnology, 156, 195–234. 10.1007/10_2016_5

Serna-García, R., Zamorano-López, N., Seco, A., & Bouzas, A. (2020). Co-digestion of harvested microalgae and primary sludge in a mesophilic anaerobic membrane bioreactor (AnMBR): Methane potential and microbial diversity. Bioresource Technology, 298, 122521. 10.1016/j.biortech.2019.122521

Shi, Z., Zhang, L [Liang], Yuan, H., Li, X [Xiujin], Chang, Y., & Zuo, X. (2021). Oyster shells improve anaerobic dark fermentation performances of food waste: Hydrogen production, acidification performances, and microbial community characteristics. Bioresource Technology, 335, 125268. 10.1016/j.biortech.2021.125268

Söhngen, C., Bunk, B., Podstawka, A., Gleim, D., & Overmann, J. (2014). Bacdive--the Bacterial Diversity Metadatabase. Nucleic Acids Research, 42(Database issue), D592-9. 10.1093/nar/gkt1058

St-Pierre, B., & Wright, A. D. G. (2014). Comparative metagenomic analysis of bacterial populations in three full-scale mesophilic anaerobic manure digesters. Applied Microbiology and Biotechnology, 98(6), 2709–2717. 10.1007/s00253-013-5220-3

Sundberg, C., Al-Soud, W. A., Larsson, M., Alm, E., Yekta, S. S., Svensson, B. H., Søren-sen, S. J., & Karlsson, A. (2013). 454 pyrosequencing analyses of bacterial and archaeal richness in 21 full-scale biogas digesters. FEMS Microbiology Ecology, 85(3), 612–626. 10.1111/1574-6941.12148

Świątczak, P., Cydzik-Kwiatkowska, A., & Zielińska, M. (2019). Treatment of the liquid phase of digestate from a biogas plant for water reuse. Bioresource Technology, 276, 226–235. 10.1016/j.biortech.2018.12.077

Theuerl, S., Klang, J., Heiermann, M., & Vrieze, J. de (2018). Marker microbiome clusters are determined by operational parameters and specific key taxa combinations in anaerobic digestion. Bioresource Technology, 263, 128–135. 10.1016/j.biortech.2018.04.111

Theuerl, S., Klang, J., Hülsemann, B., Mächtig, T., & Hassa, J. (2020). Microbiome Diversity and Community-Level Change Points within Manure-based small Biogas Plants. Microorganisms, 8(8). 10.3390/microorganisms8081169

Tomazetto, G., Hahnke, S., Wibberg, D., Pühler, A., Klocke, M., & Schlüter, A. (2018). Proteiniphilum saccharofermentans str. M3/6t isolated from a laboratory biogas reactor is versa-tile in polysaccharide and oligopeptide utilization as deduced from genome-based metabolic reconstructions. Biotechnology Reports (Amsterdam, Netherlands), 18, e00254. 10.1016/j.btre.2018.e00254

Tufaner, F., & Avşar, Y. (2016). Effects of co-substrate on biogas production from cattle manure: a review. International Journal of Environmental Science & Technology, 13(9), 2303– 2312. 10.1007/s13762-016-1069-1

VDI-Gesellschaft Energie und Umwelt (2016). Fermentation of Organic Materials—Characterization of the Substrate, Sampling, Collection of Material Data, Fermentation Tests (Norm VDI 4630). Berlin, Germany. Beuth Verlag GmbH.

Vrieze, J. de, Raport, L., Roume, H., Vilchez-Vargas, R., Jáuregui, R., Pieper, D. H., & Boon, N. (2016). The full-scale anaerobic digestion microbiome is represented by specific marker populations. Water Research, 104, 101–110. 10.1016/j.watres.2016.08.008

Vrieze, J. de, Saunders, A. M., He, Y., Fang, J., Nielsen, P. H [Per Halkjaer], Verstraete, W., & Boon, N. (2015). Ammonia and temperature determine potential clustering in the anaerobic digestion microbiome. Water Research, 75, 312–323. 10.1016/j.wa-tres.2015.02.025

Wang, C [Chen], Liu, J [Jieyi], Xu, X., & Zhu, L. (2022). Response of methanogenic granules enhanced by magnetite to ammonia stress. Water Research, 212, 118123. 10.1016/j.watres.2022.118123

Wang, H., Qu, Y., Da Li, Ambuchi, J. J., He, W., Zhou, X., Liu, J [Jia], & Feng, Y. (2016). Cascade degradation of organic matters in brewery wastewater using a continuous stirred microbial electrochemical reactor and analysis of microbial communities. Scientific Reports, 6, 27023. 10.1038/srep27023

Wang, Y [Yuanyuan], Zhang, Y [Yanlin], Wang, J., & Meng, L. (2009). Effects of volatile fatty acid concentrations on methane yield and methanogenic bacteria. Biomass and Bioenergy, 33(5), 848–853. 10.1016/j.biombioe.2009.01.007

Weiland, P. (2010). Biogas production: Current state and perspectives. Applied Microbiology and Biotechnology, 85(4), 849–860. 10.1007/s00253-009-2246-7

Westerholm, M., Moestedt, J., & Schnürer, A. (2016). Biogas production through syntrophic acetate oxidation and deliberate operating strategies for improved digester performance. Applied Energy, 179, 124–135. 10.1016/j.apenergy.2016.06.061

Wu, X., Yao, W., Zhu, J., & Miller, C. (2010). Biogas and CH(4) productivity by co-digesting swine manure with three crop residues as an external carbon source. Bioresource Technology, 101(11), 4042–4047. 10.1016/j.biortech.2010.01.052

Wu, Y., Wang, C [Cuiping], Liu, X., Ma, H., Wu, J., Zuo, J., & Wang, K. (2016). A new method of two-phase anaerobic digestion for fruit and vegetable waste treatment. Bioresource Technology, 211, 16–23. 10.1016/j.biortech.2016.03.050

Wu, Z., Nguyen, D., Lam, T. Y. C., Zhuang, H., Shrestha, S., Raskin, L., Khanal, S. K., & Lee, P. H. (2021). Synergistic association between cytochrome bd-encoded Proteiniphilum and reactive oxygen species (ROS)-scavenging methanogens in microaerobic-anaerobic digestion of lignocellulosic biomass. Water Research, 190, 116721. 10.1016/j.watres.2020.116721

Xu, H., Wang, K., Zhang, X., Gong, H., Xia, Y., & Holmes, D. E. (2020). Application of in-situ H2-assisted biogas upgrading in high-rate anaerobic wastewater treatment. Bioresource Technology, 299, 122598. 10.1016/j.biortech.2019.122598

Xu, R., Zhang, K., Liu, P., Khan, A., Xiong, J., Tian, F., & Li, X [Xiangkai] (2018). A critical review on the interaction of substrate nutrient balance and microbial community structure and function in anaerobic co-digestion. Bioresource Technology, 247, 1119–1127. 10.1016/j.biortech.2017.09.095

Yenigün, O., & Demirel, B. (2013). Ammonia inhibition in anaerobic digestion: A review. Process Biochemistry, 48(5-6), 901–911. 10.1016/j.procbio.2013.04.012

Yu, D., Zhang, J [Junya], Chulu, B., Yang, M., Nopens, I., & Wei, Y. (2020). Ammonia stress decreased biomarker genes of acetoclastic methanogenesis and second peak of production rates during anaerobic digestion of swine manure. Bioresource Technology, 317, 124012. 10.1016/j.biortech.2020.124012

Zhang, J [Junyu], Gao, Q., Zhang, Q [Qiuting], Wang, T., Yue, H., Wu, L., Shi, J [Jason], Qin, Z., Zhou, J., Zuo, J., & Yang, Y. (2017). Bacteriophage-prokaryote dynamics and interaction within anaerobic digestion processes across time and space. Microbiome, 5(1), 57. 10.1186/s40168-017-0272-8

Zhang, L [Liguo], Ban, Q., & Li, J. (2018). Microbial community dynamics at high organic loading rates revealed by pyrosequencing during sugar refinery wastewater treatment in a UASB reactor. Frontiers of Environmental Science & Engineering, 12(4). 10.1007/s11783-018-1045-8

Zhang, Q [Quanguo], Hu, J., & Lee, D.J. (2016). Biogas from anaerobic digestion processes: Research updates. Renewable Energy, 98, 108–119. 10.1016/j.renene.2016.02.029

Zhang, S., Xiao, M., Liang, C., Chui, C., Wang, N., Shi, J [Jiping], & Liu, L. (2022). Multivariate insights into enhanced biogas production in thermophilic dry anaerobic co-digestion of food waste with kitchen waste or garden waste: Process properties, microbial communities and metagenomic analyses. Bioresource Technology, 361, 127684. 10.1016/j.biortech.2022.127684

Zheng, D., Wang, H. Z., Gou, M., Nobu, M. K., Narihiro, T., Hu, B., Nie, Y., & Tang, Y. Q. (2019). Identification of novel potential acetate-oxidizing bacteria in thermophilic methanogenic chemostats by DNA stable isotope probing. Applied Microbiology and Biotechnology, 103(20), 8631–8645. 10.1007/s00253-019-10078-9

Zheng, Z., Cai, Y., Zhang, Y [Yue], Zhao, Y., Gao, Y., Cui, Z., Hu, Y., & Wang, X [Xiaofen] (2021). The effects of C/N (10-25) on the relationship of substrates, metabolites, and microorganisms in “inhibited steady-state” of anaerobic digestion. Water Research, 188, 116466. 10.1016/j.watres.2020.116466

